# Splicing factor SRSF1 is a pH-stat to restore nucleolar integrity and function

**DOI:** 10.1101/2025.04.12.648491

**Authors:** Suibin Ma, Jierui Guo, Xiang Zhan, Shuo Yang, Chaoqun Huang, Bo Wang

**Author notes:** **These authors contributed equally** (Suibin Ma, Jierui Guo). **Correspondence:** Bo Wang, State Key Laboratory of Cellular Stress Biology, School of Life Sciences, Faculty of Medicine and Life Sciences, Xiamen University, Xiamen, China 361102, China.

## Abstract

Nucleolus is a multiphase biomolecular assembly containing three distinct subcompartments. Evidence is just emerging that nucleolar proteins with characteristic electrochemical properties condense to generate a pH gradient, which serves as a key driving force to organize the nucleolar sub-phases. Given the indispensable functionality of nucleolus in most, if not all, of the cellular activities, it is vital for a cell to sense potential fluctuation in the nucleolar pH and subsequently maintain the pH homeostasis. The mechanisms how a cell achieve this task remains poorly decoded. Here, we show that splicing factor SRSF1 is shuttled from nuclear speckles (NSs) to the nucleolus in times of stress that interrogates the nucleolar pH. SRSF1 nucleolar localization is reliant on an acidic patch and molecular interactions with a nucleolar-resident protein, DDX18. Loss of SRSF1 impedes pH homeostasis in the nucleolus, and subsequently obstructs the restoration of the nucleolar multiphase and function. The arginine residues in the RS (arginine/serine rich) domain of SRSF1, endowed by high isoelectric point (pI), directly alkalize the nucleolar microenvironment. Interestingly, synthetic arginine-rich dipeptides derivative of SRSF1 RS domain safeguard nucleolus from pH and functional disturbance. Our findings uncover unprecedented mechanistic insights into nucleolar pH-sensing and regulation.

## Introduction

Nucleolus is thus far the largest membrane-less organelle that serves as the primary hub for ribosomal biogenesis, including ribosomal rRNA transcription, processing, and ribosomal assembly^1,2^. It consists of three co-existing immiscible sub-phases, namely the granular component (GC), dense fibrillar component (DFC), and fibrillar center (FC), each one of which fulfills specific steps in the process of ribosomal biogenesis. The organization and maintenance of nucleolus multiphase are dynamically regulated under various physiopathological conditions. Deciphering the molecular mechanisms underlying the organization, maintenance, and remodeling of nucleolus is of great clinical significance as disturbed dynamics and functionality of nucleolus have been shown to tightly associate with many diseases, including cancer and neurodegeneration^3^.

The extant data suggests an indispensable role of biomolecular phase separation in establishing the nucleolar multilayered structure. Mounting evidence demonstrates that the summation of protein-protein, protein-RNA, RNA-RNA interactions drive multicomponent phase separation in the nucleolus^1,4^. The nucleolus is characterized with striking thermodynamics. In addition to the constant molecular exchange with the nucleoplasm, the multiphase architecture can readily undergo cycles of destruction and remodeling^5,6^. For example, inside out nucleolar structure known as nucleolar caps can form as a response to a number of stimulations^7-9^. The context-dependent dynamic organization, maintenance and remodeling of the nucleolus multiphase is a fundamental biological problem, but remain partially understood. Recent studies have revealed that each nucleolar sub-phase has a specific pH, and the net charge of the nucleolus proteins plays a critical role in establishing a pH gradient across the nucleolar layers^10^. This internal pH gradient is sharply distinct from that detected cross the membrane of classical organelles (e.i. mitochondria), the establishment of which is not reliant on ATP-fueled selective proton pumps, but rather the different electrochemical potential across the interface of biomolecular condensates. While pH is emerging as a key chemical determinant for nucleolar organization, the regulation of nucleolar pH under different physiopathological conditions are yet to be systematically elucidated.

The serine/arginine-rich splicing factor (SRSF) protein family is a relatively well-studied group of NS-resident splicing factors participating in a plethora of cellular processes^11^. It is best studied in the context of tumorigenesis^12^. Here, we show that several members of the SRSF1 family, including SRSF1, relocate from NSs to the nucleolus when the nucleolus becomes alkalized. The pH-dependent nucleolar localization of SRSF1 is reliant on molecular interactions with a nucleolar-resident protein, DDX18, via an acidic charge block of SRSF1. Loss of SRSF1 impedes the establishment of alkaline nucleolar pH, and subsequently obstructs the restoration of the nucleolar sub-phases and function in rRNA processing. Arginine in the RS (arginine/serine rich) domain of SRSF1 is solely responsible for modulating the nucleolar pH due to its high isoelectric point. Interestingly, synthetic arginine-rich dipeptides, derivative of the SRSF1 RS domain, protect the nucleolus from pH and functional disturbance.

## Results

### SRSF1 is recruited into nucleolus under specific stressed conditions

The SR family members are nuclear speckle (NS)-resident constituents that are well known for their exclusively participation in mRNA alternative splicing^11^. Given the mounting evidence suggesting that proteins are dynamically relocated among membrane-less cellular compartments under different pathophysiological conditions, we sought to determine if the SRSF family members are capable of doing so. Towards this end, we tagged the endogenous loci of the archetypal member of the SRSF family, *SRSF1* and *SRSF2*, with GFP coding sequence via CRISPR-based knock-in (KI) technique in HeLa cells (**Figure S1A, S1B**). We then subjected these cells to an array of stressors, such as oxidative stress, UV stress, etc. We readily observed that many of these stressors drastically altered the spatial distribution of SRSF1 and SRSF2 in cells (**Figure S1C**). For instance, oxidative stress imposed by NaAsO_2_ exposure led to the emergence of cytoplasmic SRSF1 and SRSF2 foci that were suspected to be stress granules. Consistent with prior studies, heat shock triggered the formation of nuclear foci of SRSF1^13,14^, but not SRSF2 (**Figure S1C**). Interestingly, several treatments, including actinomycin D (AMD), UV, and H_2_O_2_ resulted in marked translocation of SRSF1, and to a lesser extent, SRSF2 into the nucleolus (**Figure S1C**). Although the observation that SRSF1 relocates to the nucleolus in response to specific stressors has been made decades ago, the underlying regulation and function still remain mysterious^7^. We therefore focused on this for the subsequence study. We also noticed that the stress-induced nucleolar localization of SRSF1 was more pronounced in U-2 OS cells, and hence primarily used this cell type throughout the investigation.

We first attempted to survey the dynamics of SRSF1 in the course of UV treatment (30 J/m^2^) and recovery. We found that the emergence of SRSF1 in the nucleolus gradually increased in the process of recovery from UV treatment and peaked at 7 h (**Figure 1A, 1B**), whereas the overall intracellular levels of SRSF1 remains stable (**Figure S1D, S1E**). This UV-dependent translocation of SRSF1 into the nucleolus was independently detected in several unrelated cell types such as 293T, THP-1, and HCT 116 cells (**Figure S1F-H**). The cellular distribution of other NS-associated proteins such as SRRM2 remained unaltered by UV stress (**Figure S1I**). SRSF1 consist of two RNA recognition motifs (RRMs) and a C-terminal arginine/serine rich domain (RS) domain (**Figure 1C**). We generated SRSF1 truncations lacking each individual domain to understand the structural basis of SRSF1’s localization to the nucleolus (**Figure 1D**). We found that the RRM, and specifically RRM1 region, but not the RS domain of SRSF1, was indispensable for its nucleolar enrichment (**Figure 1E, 1F**).

**Figure 1.**
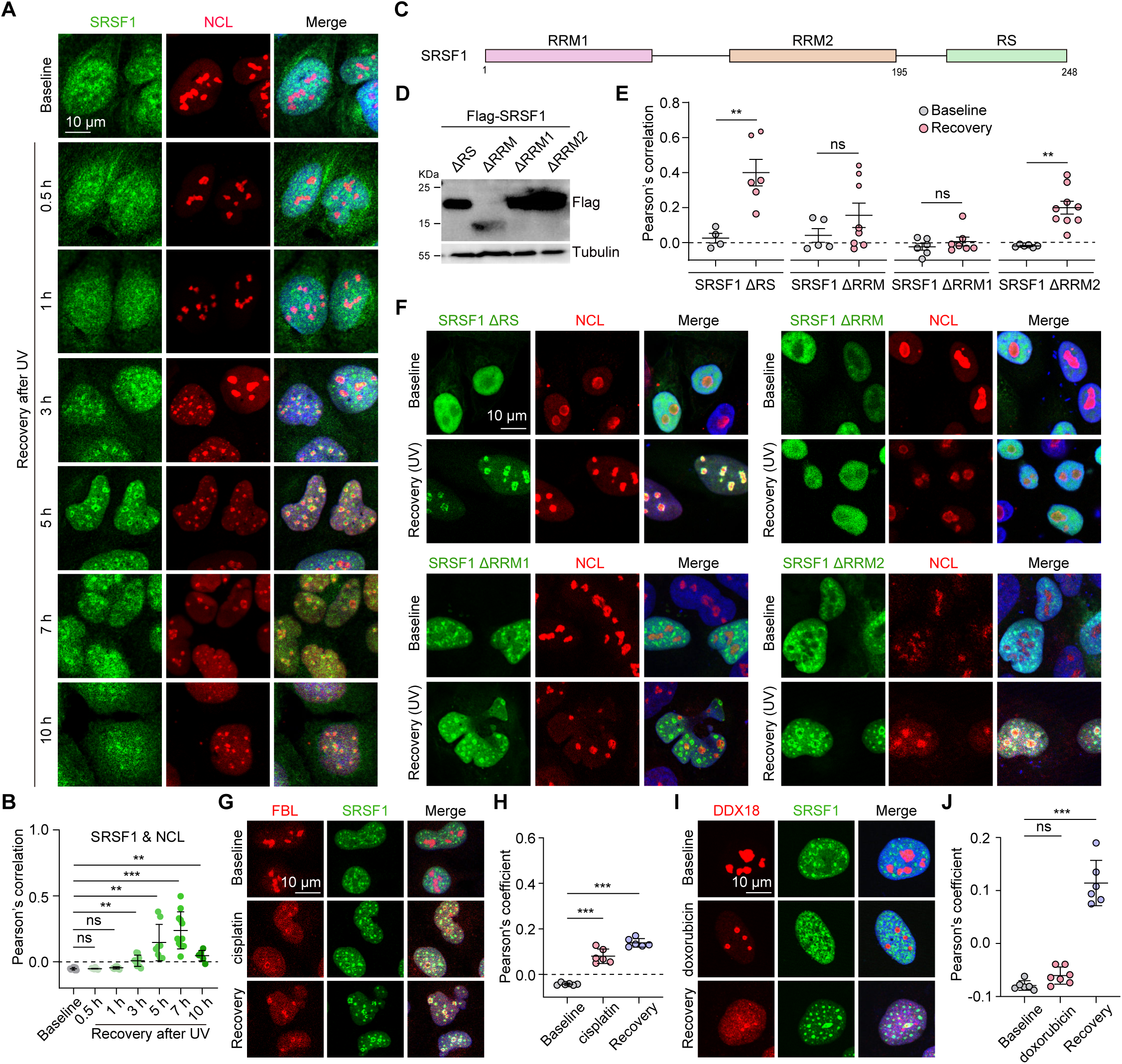
SRSF1 is recruited into the nucleolus under specific stressed conditions. (**A**, **B**) SRSF1-GFP KI U-2 OS cells were treated with UV and allowed to recover for the indicated time. The cells were further processed for immunostaining using the indicated antibodies (A). Pearson’s correlation of SRSF1-GFP and NCL were determined (B). (**C**) Schematic illustration of SRSF1 domains. (**D**) The indicated constructs were expressed in 293T cells, and the proteins were detected by immunoblotting. (**E**, **F**) WT U-2 OS cells transfected with the indicated SRSF1 mutants were stressed by UV and allowed to recover for 7 h. The cells were immunostained with the indicated antibodies (E). Pearson’s correlation of SRSF1 mutants and NCL were quantified (F). (**G**-**J**) WT U-2 OS cells were treated with either cisplatin (G, H) or doxorubicin (I, J) and recovered in normal growth medium followed by immunostaining using the indicated antibodies. Colocalization of SRSF1 with nucleolus markers were quantified (I, J). For (**B**), (**H**) and (**J**), data are shown as mean ± SD. ns, not significant; **p < 0.01; ***p < 0.001 by one-way ANOVA. For (**E**), data are shown as mean ± SD. ns, not significant; **p < 0.01 by unpaired Student’s t test.

Previous studies have shown that rRNA is integral to the establishment of nucleolar multilayers and that inhibition of rRNA biogenesis results in nucleolar remodeling, which gives rise to inside out nucleolar structure known as nucleolar cap^8,10,15,16^. We hypothesized that SRSF1’s translocation into the nucleolus is coupled with rRNA inhibition. Indeed, treating the cells with two of the most commonly used chemotherapeutic chemicals, cisplatin and doxorubicin, both of which halt rRNA biogenesis, elicited pronounced formation of nucleolar caps. Interestingly, SRSF1 were evidently enriched in the nucleolar caps caused by either cisplatin or doxorubicin treatment (**Figure 1G-J**). Moreover, UV exposure also led to dramatically reduced rRNA production (**Figure S1J-M**), which was associated with nucleolar cap formation and SRSF1 enrichment, and exposure of high dose AMD (1 µM) to inhibit the activity of RNA polymerase I (Pol I) resulted in similar phenomenon^7,8^. Together, we conclude that SRSF1 recruitment into the nucleolar caps is probably coupled with stressors specifically interrogating rRNA biogenesis.

### SRSF1 localization to the nucleolar caps is dependent on the molecular interactions with DDX18 through an acidic charge block

To understand the biochemical basis of SRSF1’s nucleolar localization, we first generated Flag-tagged SRSF family members and surveyed if the stress-dependent nucleolar recruitment is universal to the SRSFs (**Figure S2A, S2B**). We found that, in addition to SRSF1, several other members of this family including SRSF5-9, and 12 were selectively enriched in the nucleolar caps upon cellular recovery from UV treatment (**Figure S2C, S2D**). Among these SRSFs, SRSF5, 6, and 9 share similar structural domains with SRSF1, harboring two RRMs and a C-terminal RS domain (**Figure S2A**). Given that RRM1 mediates SRSF1’s localization to the nucleolus, we therefore aligned the linear sequence of SRSF1 RRM1 with those of SRSF5, 6, and 9. Interestingly, these RRM1s share considerable sequence similarity (**Figure 2A**). A cluster of negatively charged amino acids (referred to as acidic charge block) caught our attention (**Figure 2B**) because such sequence features were reported to be overrepresented in the nucleolar proteome^10^. Therefore, we speculated that the acidic charge block of SRSF1 RRM1 region confers a nucleolar localization signal. We mutated several acidic residues in this region to alanine (referred to as SRSF1^D/E-A^), which greatly diminished the negative charge and significantly reshaped the charge landscape of SRSF1 (**Figure 2A, 2B**). Indeed, D/E-A mutant exhibited dramatically compromised UV-dependent nucleolar localization of SRSF1 (**Figure 2C, 2D**).

**Figure 2.**
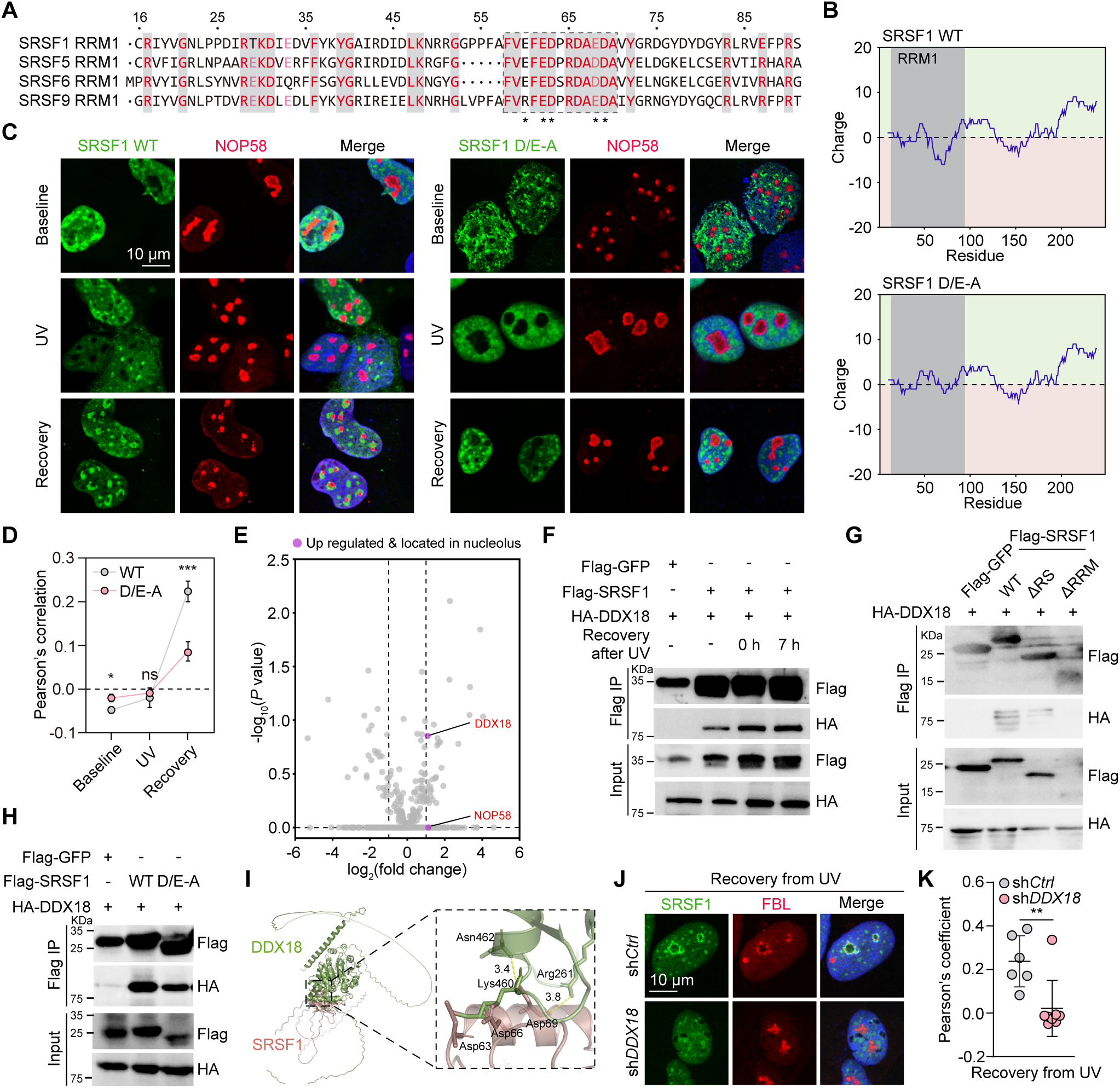
SRSF1 localization to nucleolus is dependent on molecular interaction with DDX18 through an acidic charge block. (**A**) Sequence alignment of the RRM1 regions of SRSF1, SRSF5, SRSF6, and SRSF9. (**B**) Charge plots of SRSF1 WT and D/E-A mutant. (**C**, **D**) WT U-2 OS cells transfected with SRSF1 WT or D/E-A mutant were stressed by UV and allowed to recover for 7 h. The cells were immunostained with the indicated antibodies (C). Pearson’s correlation of SRSF1 WT or D/E-A mutant and NOP58 were quantified (D). (**E**) Volcano plot of enriched protein targets from SRSF1 immunoprecipitation-mass spectrometry experiment. The targets that were both enriched in SRSF1 immunoprecipitation and located in the nucleolus were highlighted. (**F**) 293T cells transfected with the indicated constructs were stressed with UV and recovered for 7 h before being subjected to Flag immunoprecipitation. (**G**, **H**) 293T cells were transfected with the indicated constructs and subjected to Flag immunoprecipitation. (**I**) Structural prediction of SRSF1 in complex with DDX18. Key residues mediating SRSF1-DDX18 interaction were highlighted. (**J**, **K**) WT U-2 OS cells infected with shRNA against control (*Ctr*) or *DDX18* were stressed with UV followed by recovery in normal growth condition for 7 h. The cells were then processed for immunostaining with the indicated antibodies (J). Pearson’s correlation of SRSF1 and FBL were quantified (K). For (**D**), and (**K**), data are shown as mean ± SD. ns, not significant; *p < 0.05; **p < 0.01; ***p < 0.001 by unpaired Student’s t test.

To understand the molecular basis of SRSF’s nucleolar localization, we immunoprecipitated SRSF1 from cells under normal growth condition and UV-stimulated condition, and performed mass spectrometry analysis. We identified two potentially interesting targets, DDX18 and NOP58, which not only were nucleolar constituents, but also showed enhanced interactions with SRSF1 upon UV treatment (**Figure 2E**). Targeted immunoprecipitations validated the SRSF1-DDX18, and SRSF1-NOP58 interactions, both of which were evidently increased by UV treatment (**Figure 2F, and S2E**). The RRM domain was responsible for SRSF1’s interaction with both DDX18 and NOP58 (**Figure 2G, and S2F**). Given the indispensability of the acidic charge block of SRSF1 RRM1 region for the nucleolar enrichment, we sought to determine if the acidic charge block modulates the molecular interactions of SRSF1 with DDX18 and SRSF1. Disrupting the acidic charge block prominently reduced the molecular interaction of SRSF1 and DDX18 (**Figure 2H**). Molecular docking revealed an interesting positively charged pocket of DDX18 readily to engage in electrostatic attraction with the negative charge block of SRSF1 (**Figure 2I**). In contrast, SRSF1-NOP58 binding was not significantly blunted by the D/E-A mutation (**Figure S2G**). These data suggest that SRSF1-DDX18 interaction through the acidic charge block might promote SRSF1 recruitment into the nucleolus. Indeed, depleting the endogenous DDX18 substantially impaired stress-dependent nucleolar localization of SRSF1 (**Figure 2J, 2K, and S2H**). Consistently, silencing NOP58 had no appreciable effect on SRSF1 nucleolar recruitment (**Figure S2I-K**). We therefore conclude that enrichment of SRSF1 in the nucleolar caps is fulfilled through the molecular interactions with DDX18 via electrostatic attraction.

### SRSF1 senses and regulates nucleolar pH

Recent evidence suggests that nucleolar proteins are uniquely enriched with charged residues, which set up a pH gradient between the nucleolus and nucleoplasm, and RNA molecules, including nucleolar RNA and rRNA, contribute to nucleolar acidification via complex coacervation^10,15,16^. Given our above-described observation that SRSF1 recruitment into the nucleolar caps is coupled with the inhibition of rRNA biogenesis, we hypothesized that the inhibited rRNA biogenesis alters nucleolar pH, which subsequently resulted in SRSF1 shuttling into the nucleolus. In this case, SRSF1 senses the fluctuation in the nucleolar pH. To test it, we cultured U-2 OS cells in medium with varying pH. While both acidic and alkaline culture conditions notably disrupted the nucleolar multiphase organization, only the later triggered the marked translocation of SRSF1 from NSs to the nucleoli (**Figure 3A, 3B**). This was similarly observed in MDA-MB-231 and HCT 116 cells (**Figure S3A-D**).

**Figure 3.**
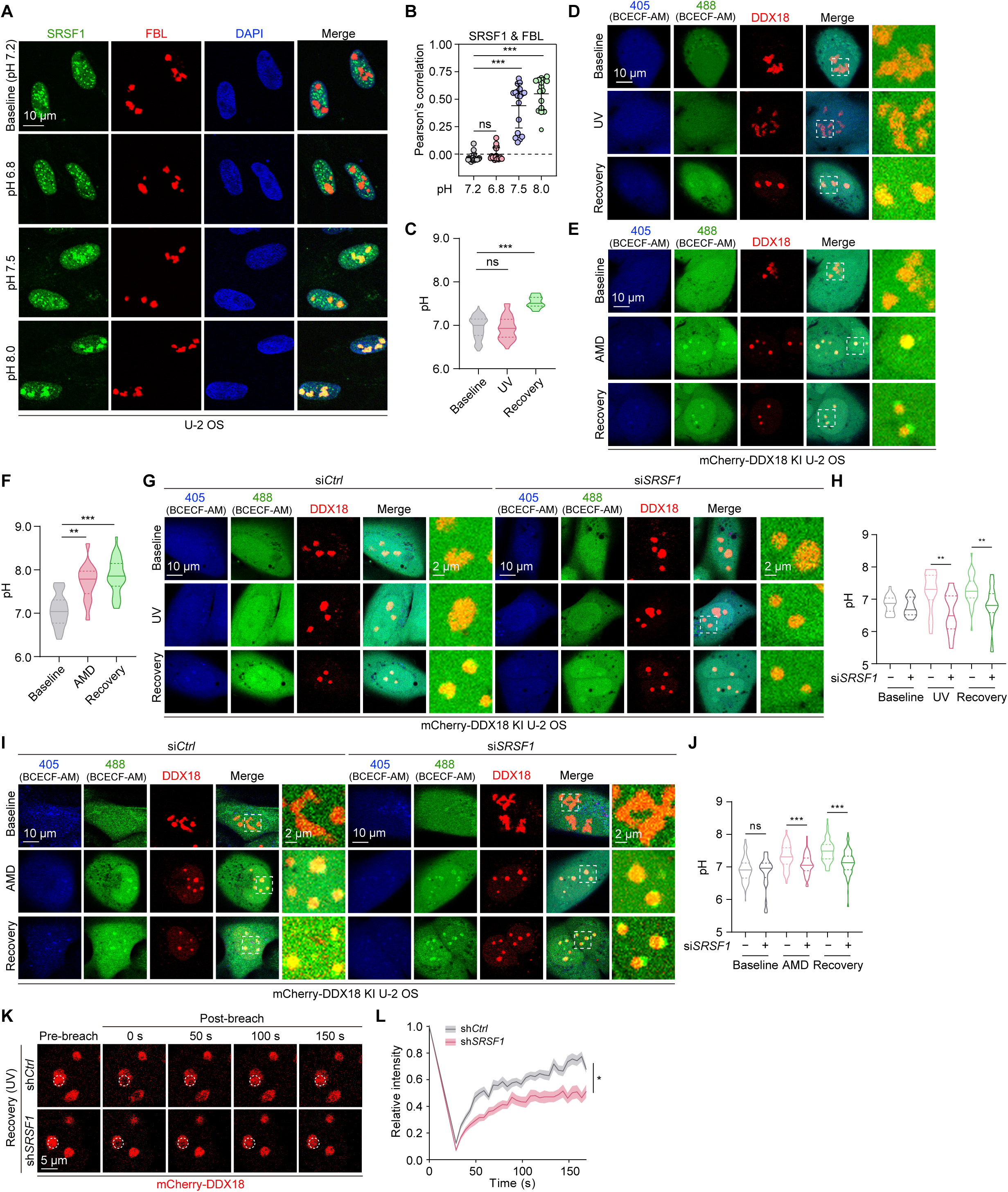
SRSF1 senses and maintains nucleolar pH. (**A**, **B**) WT U-2 OS cells were cultured at different pH. The cells were immunostained with the indicated antibodies. Colocalization of SRSF1 with FBL was quantitatively assessed by Pearson’s correlation. Data are shown as mean ± SD. ns, not significant; **p < 0.01 by one-way ANOVA. (**C**-**F**) Live cell imaging of mCherry-DDX18 KI U-2 OS being treated UV (C, D), or doxorubicin (E, F) and recovered in normal growth condition with a pH indicator BCECF-AM. pH was quantified by calculating the ratio of the fluorescence intensity of 488/405 nm in the nucleoli. ns, not significant; **p < 0.01; ***p < 0.001 by one-way ANOVA. (**G**-**J**) mCherry-DDX18 KI U-2 OS transfected with si*Ctrl* or si*SRSF1* were treated with either UV (G, H), or AMD (I, J), and recovered in normal growth condition. Live cell imaging was performed to measure nucleolar pH. ns, not significant; **p < 0.01; ***p < 0.001 by one-way ANOVA. (**K**, **L**) WT U-2 OS cells transduced with shRNA against *Ctrl* or *SRSF1* were treated with UV and recovered for 7 h. The cells were subjected to FRAP. The fluorescence intensity of mCherry-DDX18 was monitored over time and quantified (L). Data are shown as mean ± SD. n = 21. *p < 0.05 by two-way ANOVA.

Given that pH varies within the nucleolar sub-phases, we tagged the endogenous loci of *DDX18* and *FBL*, which encoded proteins residing in the nucleolar GC and DFC, respectively, with mCherry coding sequence in U-2 OS cells to directly quantitate the stress-dependent pH regulation in the nucleolus (**Figure S3E, S3F**). We performed live cell imaging by utilizing a membrane-permeable dye BCECF-AM to quantitatively measure the nucleolar pH^17,18^. This fluorescence dye shows pH-sensitive emission intensity when the dye is excited at ∼490 nm and pH-insensitive emission intensity when excited ∼440 nm. Therefore, intracellular pH can be quantitatively determined by calculating the emission ratio. Interestingly, the DDX18^+^ nucleolar caps formed during the cellular recovery from UV treatment were dramatically alkaline (**Figure 3C, 3D**). Treating cells with other stressors including AMD, cisplatin, and doxorubicin that impeded rRNA biogenesis similarly resulted in the formation of highly alkaline DDX18^+^ nucleolar caps (**Figure 3E, 3F, and S3G-J**). Importantly, no significant pH alteration within the Fibrillarin/FBL^+^ subcompartment was induced by any of the above stressors (**Figure S3K-R**).

To determine if SRSF1 regulates the nucleolar pH, we investigated the potential pH changes due to SRSF1 interference. We found that although SRSF1 silencing did not abolish the formation of nucleolar caps incited by UV treatment, these structures were significantly more acidic in the absence of SRSF1 (**Figure 3G, 3H**). In a similar fashion, when SRSF1 was not present in the nucleolar caps induced by AMD, the compartments became remarkably less alkaline (**Figure 3I, 3J**). Together, our data implicate a nucleolar deacidification-dependent recruitment of SRSF1, which further contributes to the nucleolar deacidification. We reasoned that the difference in pH between the nucleolar caps and the surrounding milieu generates interphase electric potentials to power molecular diffusion in the nucleolar caps. Indeed, SRSF1-deficiency caused a dramatically reduced recovery rate after fluorescence photobleaching, suggesting that SRSF1-maintained nucleolar pH was essential for accelerating molecular motility (**Figure 3K, 3L**).

### SRSF1 maintains nucleolar integrity and function in times of stress

To determine the functional role of SRSF1 in managing the nucleolar stress, we selective silenced SRSF1 with either shRNA or siRNA and systematically analyzed the subsequent impact on nucleolar organization and function. Both UV and AMD treatment led to the formation of inside out nucleolar caps with dislocated DFC to reside outside of the GC layer. The characteristic multilayer organization gradually restored during cellular recovery from the stressed conditions. In cells with depleted SRSF1, re-establishment of nucleolar multiphases during recovery from either UV irradiation or AMD treatment was dramatically impeded, associated with significantly reduced nucleolar size (**Figure 4A-F**). We examined the distribution of GC-resident protein NPM1, finding that the partition of NPM1 into the GC phase was greatly diminished in sh*SRSF1* transduced cells compared to the scramble shRNA (**Figure S4A, S4B**). Next, we utilized an EU incorporation assay, a fluorescence in situ hybridization (FISH) probe for 5’-ETS, and an antibody specific against mature rRNA to assess the transcriptional activity of rRNA, processing and maturation of rRNA, respectively. We found that the obstructed nucleolar restoration due to the loss of SRSF1 resulted in compromised rRNA processing and maturation (**Figure 4G-J**), but not transcription (**Figure S4C, S4D**). Given that UV irradiation causes widespread DNA damage and the nucleolus is the central hub for DNA repair, we sought to determine if SRSF1 participate in this process. We immunostained the cells with an antibody selectively against γH2AX, to visualize foci of DNA damage, and detected significantly higher levels of DNA damage in SRSF1 knockdown cells (**Figure 4K, 4L**). In a similar fashion, SRSF1 knockdown also led to dramatically hampered rRNA processing and maturation, and DNA repair in cells restored from AMD treatment (**Figure S4E-I**). We further confirmed that these compromised cellular processes in the absence of SRSF1 was due to aberrant nucleolar organization, but not the abundance of the main nucleolar constituents (**Figure S4J, S4K**).

**Figure 4.**
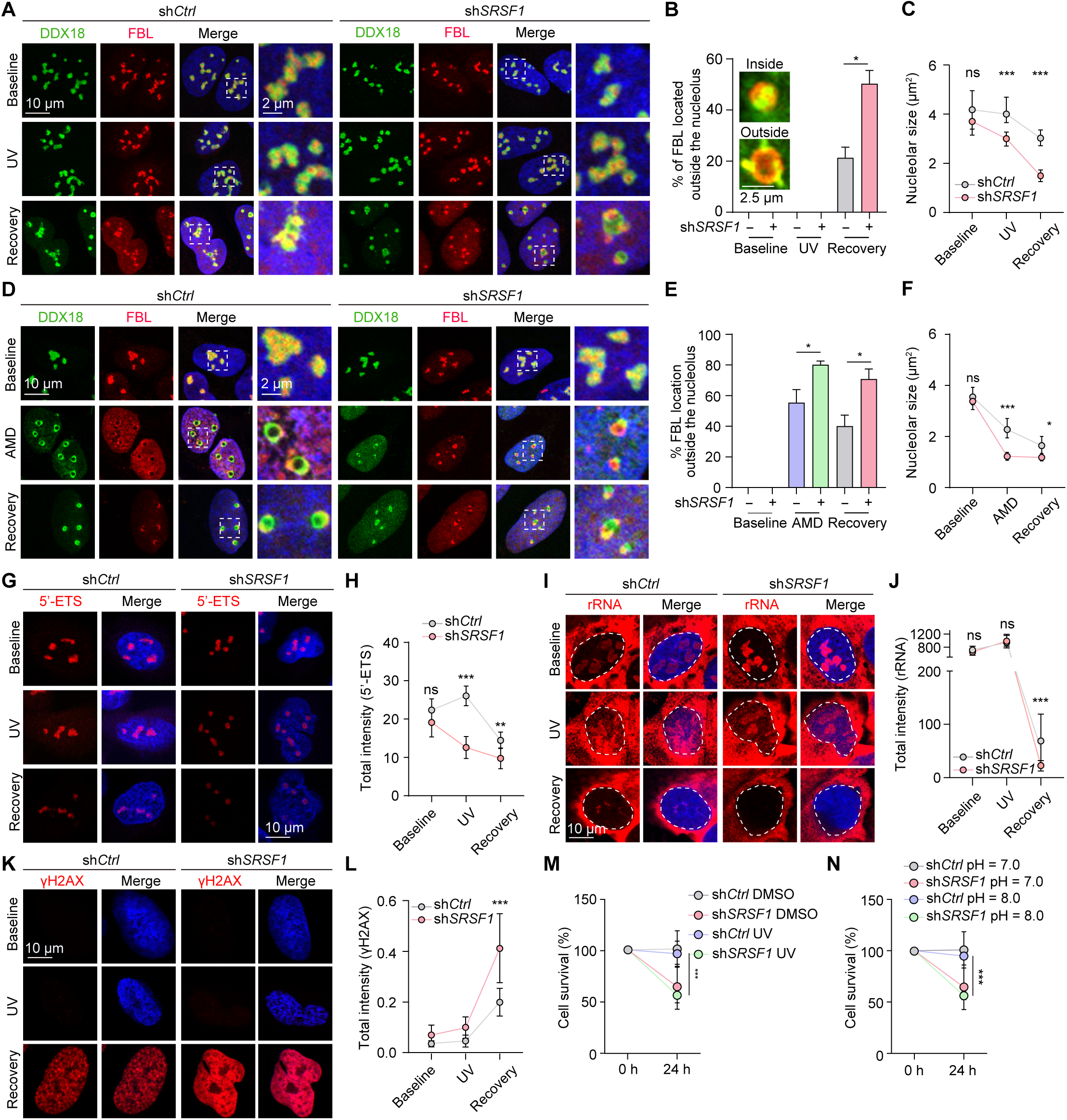
SRSF1 maintains nucleolar integrity and function in times of stress. (**A-F**) WT U-2 OS cells transduced with shRNA against *Ctrl* or *SRSF1* were treated with either UV (A-C) or AMD (D-F). The cells were immunostained with antibodies against DDX18 and FBL. Percentage of nucleoli with FBL dislocated to the periphery and the nucleolar size were quantified. (**G**-**L**) WT U-2 OS cells transduced with shRNA against *Ctrl* or *SRSF1* were treated with UV and recovered for 7 h. The cells were subjected to either FISH using a probe for 5’-ETS (G, H), immunostaining using an antibody against rRNA (I, J), or γH2AX (K, L). The total intensity of 5’-ETS (H), rRNA (J), and γH2AX (L) were quantified, respectively. (**M**, **N**) WT U-2 OS cells transduced with shRNA against *Ctrl* or *SRSF1* were treated with UV (M) or alkaline (pH = 8.0) culture medium (N). 24 h after the treatment, cell survival was determined by CCK8 assay. For (**B**), and (**E**), data are shown as mean ± SEM. n = 3 independent experiments. *p < 0.05 by unpaired Student’s t test. For (**C**), (**F**), (**H**), (**J**), and (**L**), data are shown as median ± 95% confidence interval. ns, not significant; *p < 0.05; **p < 0.01; ***p < 0.001 by unpaired Student’s t test. For (**M**), and (**N**), data are shown as mean ± SD. **p < 0.01; ***p < 0.001 by unpaired Student’s t test.

Because prolonged nucleolar dysfunction and unsolved DNA damage result in cell demise, we asked if the loss of SRSF1 would render cells more susceptible to nucleolar stress. Cell growth was considerably inhibited when UV irradiation or AMD treatment was applied. The inhibitory effect on cell growth was further heightened by SRSF1 interference (**Figure 4M, and S4M**). In a similar fashion, two other chemotherapy drugs, cisplatin and doxorubicin, which incited SRSF1 nucleolar localization (**Figure 1G-J**), also hindered cell growth. Cells became even more vulnerable to cisplatin and doxorubicin treatment when SRSF1 levels were low (**Figure S4N, S4O**). More importantly, cells were more susceptible to alkaline culture condition (pH = 8) when SRSF1 was depleted (**Figure 4N**), highlighting the importance of maintaining physiologic pH. Collectively, our data suggest that SRSF1 shuttling into the nucleolus is essential for restoring nucleolar integrity, cellular function and survival.

### SRSF1 tunes nucleolar pH and function via its alkaline RS domain

The C-terminus of SRSF1 is featured with alternating arginine and serine repeat sequence. Arginine, imparted by its high isoelectric point (pI), is a potent pH modulator. Hence, an appealing speculation is that the capability of SRSF1 in regulating nucleolar pH is directly encoded by the primary sequence of its alkaline RS domain. To test this hypothesis, we designed a number of SRSF1 mutants (**Figure 5A**). Among these, all the arginine residues in the C-terminal RS domain were substituted to either lysine (referred to as SRSF1^R-K^), which retained the net positive charge but has a lower pI (10.65) compared to SRSF1 WT (pI = 12.58), or to glutamine (referred to as SRSF1^R-Q^) that fundamentally abolished the net positive charge and decreased the pI to 6.74. Alternatively, we also replaced the serine residues with either alanine (referred to as SRSF1^S-A^) that presumably preserved both the charge landscape and pI, or aspartic acid (referred to as SRSF1^S-D^) to neutralize the positive charge and pI by introducing oppositely negative charge. Agreeing with our results that SRSF1 RRM was required for its shuttle into the nucleolar caps, mutating these C-terminal amino acids did not significantly alter its UV-dependent nucleolar localization (**Figure 5B, 5C**). Of note, SRSF1^S-A^ mutant exhibited constitutive nucleolar localization even when cells were under normal growth condition (**Figure S5A, S5B**). We then reconstituted these SRSF1 C-terminal variants into shRNA-mediated SRSF1 knockdown cells and performed the nucleolus-relevant morphological and functional analyses (**Figure S5C**). First, we investigated the associated changes in nucleolar pH by the SRSF1 mutants. As revealed by live cell imaging, WT SRSF1 robustly alkalinized the nucleolar caps. In clear contrast, the variants with lower pI including SRSF1^R-K^, SRSF1^R-Q^, and SRSF1^S-D^, failed to do so. Strikingly, reintroducing the SRSF1^S-A^ mutant resulted in the emergence of highly alkaline nucleolar caps (**Figure 5D, 5E, and S5D, S5E**).

**Figure 5.**
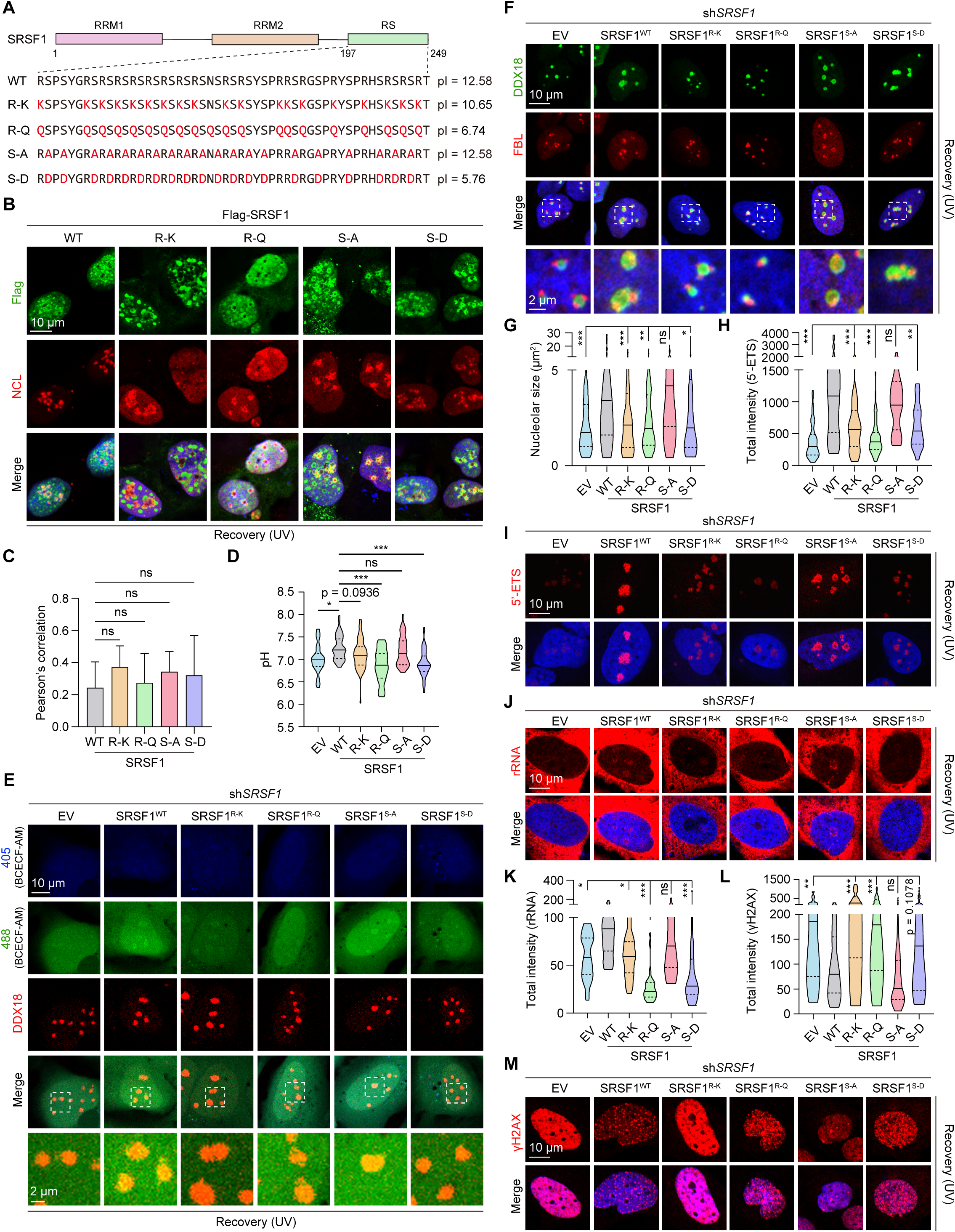
SRSF1 modulates nucleolar pH and function via its alkaline RS domain. (**A**) The linear sequences of the designed SRSF1 mutants. The corresponding pI of the C-terminal RS domains was shown on the right. (**B**, **C**) U-2 OS cells expressing the indicated SRSF1 variants were treated with UV, followed by recovery in normal growth condition for 7 h. The cells were processed for immunostaining. Pearson’s correlation of SRSF1 and NCL were quantified (C). Data are shown as mean ± SD. (**D**, **E**) WT U-2 OS cells transduced with sh*SRSF1* were further reconstituted with shRNA resistant SRSF1 variants. These cell lines were treated with UV followed by recovery for 7 h. Nucleolar pH was measured by live cell imaging. (**F**-**G**) The indicated cell lines were treated with UV followed by recovery for 7 h. The cells were immunostained with antibodies against DDX18 and FBL. The nucleolar size was quantified (**G**). (**H**-**M**) The indicated cell lines were treated with UV and recovered for 7 h. The cells were processed for either FISH using a probe for 5’-ETS (H, I), immunostaining using an antibody against rRNA (J, K), or γH2AX (L, M). The total intensity of 5’-ETS (H), rRNA (K), and γH2AX (L) were quantified, respectively. For (**C**), (**D**), (**G**), (**H**), (**K**), and (**L**), ns, not significant; *p < 0.05; **p < 0.01; ***p < 0.001 by one-way ANOVA.

Next, we collectively assessed the consequential outcomes due to the altered nucleolar pH resulted from the expression of the SRSF1 mutants. FRAP assayed again showed that the re-introducing WT SRSF1 as well as the SRSF1^S-A^ mutant to the SRSF1-deificent cells significantly fluidized the nucleolar caps (**Figure S5F, S5G**). In contrast, the nucleolar caps showed much less fluidity when SRSF1^R-K^, SRSF1^R-Q^, or SRSF1^S-D^ was reconstituted (**Figure S5F, S5G**). In addition, expression of WT SRSF1 potently rescued the impaired nucleolar remodeling and reduced nucleolar size seen in SRSF1-deficient cells. This rescue effect was comparably achieved with the expression of SRSF1^S-A^, but not any other mutants (**Figure 5F, 5G, and S5H**). Functionally, SRSF1 WT reconstitution efficiently reversed the aberrant rRNA processing, maturation, and DNA damage response elicited by the lack of SRSF1 (**Figure 5H-M**). Consistently, SRSF1^S-A^ mutant, but not the others, was able to alleviate the defective nucleolar function, to an equivalent degree as the WT protein did (**Figure 5H-M**). Again, the rescue effects were independently consolidated with experiments performed using AMD treatment (**Figure S5I-L**). Altogether, our evidence suggests that the high pI of SRSF1, most likely after reaching a threshold value, but not the net charge *per se*, is a major determinant in tuning nucleolar pH, resilience, and functionality.

### Synthetic arginine-rich alkaline dipeptides protect the nucleolus from stress that disrupts nucleolar pH and function

Inspired by the alternating dipeptide configuration of SRSF1 RS domain, we synthesized 50 repeats of RS, KS, QS, RA, and RD dipeptides to further isolate the role of the SRSF1 RS domain in tuning the nucleolar pH, structural and functional restoration. Among these, RS_50_, KS_50_, and RA_50_ dipeptide repeats carry equivalently prominent positive charge but with distinctive pI, with KS_50_ possessing a relatively lower pI value (11.69). QS_50_ and RD_50_ dipeptides are nearly neutral (**Figure 6A**). Consistently with prior ideas, peptides with high pI, including KS_50_ and RA_50_, except RS_50_, showed constitutive colocalization with the nucleolus under baseline condition (**Figure 6B, 6C**). After being challenged with high dose AMD (1 µM for 1 h), cells expressing the RS_50_, QS_50_ and RD_50_ dipeptides barely re-established the normal nucleolar structure upon drug withdrawal for 4 h (**Figure 6D, 6E**). Strikingly, the presence of exogenous RA_50_ dipeptide drastically accelerated the growing of the nucleolar size during stress recovery (**Figure 6D, 6E**). Of note, although, the KS_50_ dipeptide also showed constitutive nucleolar localization, its capability to expedite structural remodeling of the nucleolar was much less potent than the RA_50_ dipeptide (**Figure 6D, 6E**). Agreeing with these results, exogenous RA_50_ dipeptide also facilitate rRNA processing and maturation, and DNA repair for cells recovering from harsh transcriptional inhibition, whereas the KS_50_ dipeptide only marginally accelerated these processes (**Figure 6F-6I**). Note that the NS-resident RS_50_ repeat drastically relocated rRNA into NSs for unclear mechanisms. Together, our evidence suggests that peptide pI is a key determining factor for structural and functional restoration of the nucleolus. To further test this, we generated RA dipeptides of varying repeat numbers (10, 20, 30, 40, and 50) with increasing pI values (**Figure S6A**). We found that, repeat number larger than 20 was a prerequisite for nucleolar localization (**Figure S6B, S6C**). As expected, the RA dipeptides promoted the growing of nucleolar size, and subsequently the rRNA processing and DNA repair in a repeat number-dependent manner (**Figure S6D-S6I**).

**Figure 6.**
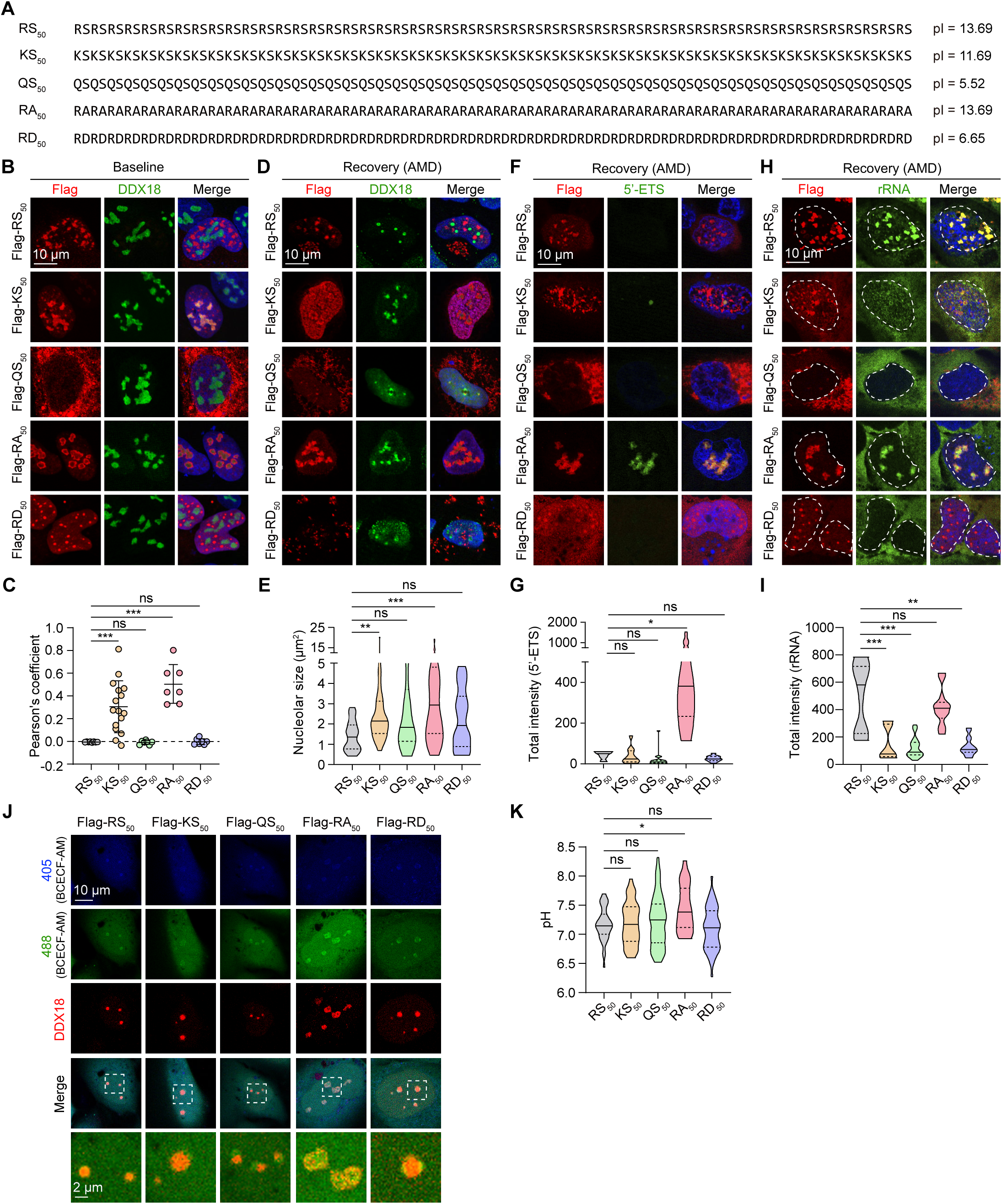
Synthetic arginine-rich alkaline dipeptide protects nucleolus from stress that disrupt nucleolar pH and function. (**A**) The linear sequences of synthetic dipeptides. The corresponding pI was shown on the right. (**B**, **C**) U-2 OS cells expressing the indicated synthetic dipeptides were immunostained with the indicated antibodies. Pearson’s correlation between DDX18 and different dipeptides was quantified (C). Data are shown as mean ± SD. ns, not significant; ***p < 0.001 by one-way ANOVA. (**D**-**I**) U-2 OS cells expressing RA dipeptide were first treated with AMD, followed by recovery in fresh medium for 4 h. The cells were further processed for FISH using a probe against 5’-ETS or immunostaining using the indicated antibodies. The nucleolar size (E), total intensity of 5’-ETS (G), and rRNA (I) were quantified, respectively. ns, not significant; *p < 0.05; **p < 0.01; ***p < 0.001 by one-way ANOVA. (**J**, **K**) Live imaging of U-2 OS cells expressing the indicated synthetic dipeptides was performed to measure nucleolar pH. ns, not significant; *p < 0.05 by one-way ANOVA.

Finally, we asked if RA dipeptide exerts protective effects toward nucleolar stress through pH modulation. Again, we employed live cell imaging to measure the possible changes in nucleolar pH due to the expression of different dipeptides. Of the tested dipeptides, only RA_50_ further alkalized the already alkaline nucleolar subcompartment resulted from AMD exposure, whereas KS_50_ dipeptide failed to do so despite of its prominent nucleolar localization (**Figure 6J, 6K**). Therefore, the pI values of synthetic arginine-rich alkaline dipeptides are positively correlated with their ability to fine-tune the nucleolar pH and resilience.

## Discussion

To the best of our knowledge, our study is the first demonstration that NS-resident SRSF1 (and likely several other members of the SRSF family) has functions most likely beyond the mRNA splicing. NSs are enriched with constituents harboring RS domains. Although the RS domain with repeat sequences has been discovered for decades, its functionality remains poorly defined^19^. We recently showed that the RS domains of NS scaffold protein SRRM2 initiate higher-order crosslinks to drive phase separation and subsequently promote NS assembly^20^. Here, we present evidence that the fundamental alkaline properties of RS domain directly encode potentials to tune the pH of nucleolar caps, which is critical for reestablishment of nucleolar multiphase structure and rRNA processing upon recovery from stress (**Figure S6J**). This pH gradient within the nucleolus is fundamentally different from the membrane potential of classical organelles such as mitochondria because its establishment does not require ATP-powered proton pumps, but rather the different electrochemical properties that are directly dictated by protein linear sequences, across the interface of biomolecular condensates. Inspired these findings, we further synthesized RA repeat dipeptides that show constitutive nucleolar localization and strikingly spare the nucleolus from stress, in a pI-dependent fashion. Interestingly, such R-rich dipeptides, PR in particular, has been shown to closely associated with the pathomechanisms of amyotrophic lateral sclerosis, a fatal motor neuron disease^21^. Patients carrying expansions of a hexanucleotide repeat GGGGCC (G_4_C_2_) in *C9ORF72*, which is the most common cause of amyotrophic lateral sclerosis, produce five distinct alternating dipeptides with one of them being PR dipeptide^21,22^. The PR dipeptide exhibits primarily nucleolar localization, is highly alkaline, but imposes significant stress on the nucleolus, leading to substantial translational inhibition and cytotoxicity^23,24^. It will be an interesting route to determine the physiochemical mechanisms underlying the very distinct biological effects of RA and PR dipeptides for future studies.

Since its first discovery by Donna Granick that transcription inhibition elicits a drastic reorganization of nucleoli, yielding the structure originally named as “’nucleolar necklace”^25,26^, nucleolar caps have been studied for nearly five decades. And yet, the exact dynamics and functionality of nuclear caps still remains largely unknown. We present quantitative evidence that the nucleolar cap is a highly alkaline cellular compartment. Of note, many of the known constituents, including SRSF1 and U1-70K, carry prominent positive charge^7^. The positively charged arginine residues of SRSF1 directly contribute to the alkaline microenvironment of the nucleolar caps. pH disturbance in this compartment due to the loss of SRSF1 results in compromised nucleolar reorganization and function. Nevertheless, the detailed functionality and physicochemical mechanisms of assembling such alkaline caps during the nucleolar stress still entail further investigation. It is likely that by maintaining a proton gradient in the nucleolar caps, it on the one hand promotes biochemical reactions such as rRNA processing, and on the other hand ensures continuous and directional flux of charged macromolecules such as rRNA or rDNA into the nucleolar interior, which in turn facilitates the nucleolar acidification. This remains an interesting hypothesis that is yet to be fully tested.

The expression of SRSF1 is upregulated in many cancer types^12^. On one hand, high levels of SRSF1 can facilitate tumorigenesis through regulation of mRNA alternative splicing. Indeed, this is well-documented by many independent investigations^27^. On the other hand, it is known for decades that acidic microenvironment is more favorable for cancer cells, likely due to the fact that cancer cells exhibit heavy reliance on ribosomal biogenesis for malignant proliferation and the nucleolus is slightly acidic as well. We observed that alkaline environment cast drastic stress upon the nucleolus. Many chemotherapy drugs, including cisplatin and doxorubicin, impose nucleolar stress and impede the ribosome production. Upregulated SRSF1 in cancer cells might promote cell survival under stressed conditions through sensing and maintaining the nucleolar pH. Therefore, it is tempting to speculate that approaches to interfere nucleolar pH, including inhibiting SRSF1, represent a new therapeutic avenue for certain cancers.

## Material and Methods

### Plasmids

Backbone vectors were linearized using DNA restriction endonucleases. Target fragments were amplified with high-fidelity DNA polymerase 2× Phanta Max Master Mix (Vazyme; P515-01). PCR primers, designed with SnapGene software, included 15 bases overlapping with the vectors. Recombinant plasmids were then assembled using the 2× MultiF Seamless Assembly Kit (Abclonal; RK21020) following the manufacturer’s instructions. Correct clones were confirmed by Sanger sequencing.

SRSF family members and their mutants were cloned into a pC3.3 vector. SRSF1 mutants for rescue assays and DPR plasmids were engineered into the pCW57.1 vector. SRSF1, DDX18 and NOP58 knockdown were achieved using shRNA against *SRSF1* (5’-TGGTCGCGACGGCTATGATTA-3’), sh*DDX18* (5’-GCACTGAGCACTGTTACTTCT-3’), and sh*NOP58* (5’-GGTGACTCCACACTTCCAACC-3’), respectively, with a pLKO.1 vector.

### Cell culture, transfection, lentivirus infection and drug treatment

U-2 OS (HTB-96), HEK293T (CRL-1573), HeLa (CCL-2), THP-1 (TIB-202), and HCT 116 (CCL-247) cell lines were purchased from (American Type Culture Collection; ATCC). U-2 OS cells were cultured in McCoy’s 5A medium (BasalMedia; L630KJ), the rest of cell lines were cultured in DMEM medium (BasalMedia; L120KJ). All media were supplemented with 10% fetal bovine serum (Moybio; S450), penicillin/streptomycin (BasalMedia; S110JV), and Glutamax (BasalMedia; S210JV) and maintained at 37°C in a cell incubator with 5% CO_2_. All the cells were routinely tested for mycoplasma contamination and were negative.

For protein overexpression experiments, transient transfection experiments were performed using Lipofectamine 2000 (Thermo Fisher; 11668019), or Neofect™ DNA transfection reagent (Neofect biotech; TF201201) according to the manufacturer’s instructions. Stable overexpression was achieved with the lentiviral pLEX plasmids. Lentiviruses were packaged in HEK293T cells. After 48 hours, viruses were collected, filtered, and used to infect target cells in the presence of 8 μg/mL polybrene (Santa Cruz Technology; sc-134220).

The transduced cells were selected after 48 h of infection with puromycin or hygromycin. All chemical drugs were diluted in DMSO or PBS as follows: 800 μM H_2_O_2_ for 1 h; 10 μM puromycin for 1 h; 0.2 M sorbitol for 2 h; 1 μM AMD (MCE; HY-17559); 100 μM cisplatin (Amjicam; ajci8058) for 6 hours; 2 μM doxorubicin (DRB) (MCE; HY-14392) for 6 h.

### RNA interference

siRNAs synthesized by GenePharma (Shanghai, China) was used: si*SRSF1*: 5’-UUGGCAGUAUUGACCUUA-3’ and 5’-UAAGGUCAAUACUGCCAA-3’. Cells were transfected with the indicated siRNAs using Lipofectamine RNAiMAX (Thermo Fisher Scientific) according to the manufacturer’s instructions and analyzed 48 h after transfection.

### Immunoblotting

Cells were lysed by lysate buffer (20 mM Tris-HCl [pH 7.5], 150 mM NaCl, 1 mM EDTA, 1 mM EDTA, 1% Triton X-100, 2.5 mM sodium pyrophosphate, 1 mM β-glycerol phosphate and a protease inhibitor cocktail). Samples were centrifuged at 10,000 rpm for 15 min at 4°C. The collected supernatant was boiled at 100°C for 10 min. The protein samples were loaded in SDS-PAGE gel, and then transferred to polyvinylidene fluoride (PVDF) membranes after electrophoresis. The membranes were blocked with 5% milk and then incubated at 4°C for overnight with the following primary antibodies: mouse anti-Flag (GNI; GNI4110-FG), mouse anti-SRSF1 (Invitrogen; 32-4500), rabbit anti-FBL (Proteintech; 16021-1-AP), rabbit anti-DDX18 (Proteintech; 66985-1-Ig), mouse anti-NPM1 (Proteintech; 10306-1-AP), rabbit anti-NOP58 (Abclonal; A4749), mouse anti-HA (Santa Cruz; sc-7392), rabbit anti-α-Tubulin (CST; 2125S), mouse anti-β-actin (Proteintech; 66009-1-Ig) and rabbit anti-NCL (Proteintech; 10556-1-AP). The membranes were then incubated with HRP-conjugated secondary antibodies (Jackson ImmunoResearch; 115-035-003, 111-035-003). Proteins were detected using a chemiluminescence detection kit (Merck Millipore; WBKLS0050).

### Immunofluorescence

Cells on the coverslip were fixed with 4% paraformaldehyde (PFA) for 10 min at room temperature (RT), blocked with 3% bovine serum albumin for 30 min RT, and incubated at 4°C for overnight with the following primary antibodies: mouse anti-SRSF1 (Invitrogen; 32-4500), rabbit anti-FBL (Proteintech; 16021-1-AP), mouse anti-FBL (Proteintech; 66985-1-Ig), rabbit anti-DDX18 (Proteintech; 66985-1-Ig), mouse anti-NPM1 (Proteintech; 10306-1-AP), rabbit anti-γH2AX (Abclonal; AP0687), rabbit anti-NOP58 (Abclonal; A4749), mouse anti-rRNA(Santa Cruz Biotechnology; SC-33678), and rabbit anti-NCL (Proteintech; 10556-1-AP). After washing three times with PBST (0.2% Triton X-100 in PBS), cells were then incubated with Alexa-Fluor-conjugated secondary antibodies (Jackson ImmunoResearch; 115-545-003, 115-585-003) for 1 h, and mounted on slides with mounting media (SouthernBiotech; 0100-01). Samples were imaged with the LSM-900 confocal microscope (ZEISS) equipped with a 63x oil objective.

### CCK8 assay

Cell proliferation assay was performed using the CCK-8 kit (GenScript CELLCOOK; L00432) according to the manufacturer’s instructions.

### Generation of KI cell lines with CRISPR/Cas9

We used the CRISPR/Cas9 system to generate U-2 OS cells with endogenous SRSF1 tagged with GFP, FBL and DDX tagged with mCherry, as well as SRSF1 and SRSF2 tagged with GFP in HeLa cells. Template sequences with 750 bp flanking homology arms for recombination were cloned into a pUC19 vector, while gRNAs were cloned into a PX459 vector. We co-transfected U-2OS cells with the vectors pUC19 and PX459 vectors containing the corresponding gRNAs. gRNA was designed using the online tool: the Optimized CRISPR Design (https://portals.broadinstitute.org/gppx/crispick/public). The guide sequences for *SRSF1*, *SRSF2*, *FBL*, and *DDX18* knock-in were 5′-GTCATAGCAGATCTCGCTCT-3′, 5′-CCCAAGTCTCCTGAAGAGGA-3′, 5′-GTTGTGAACGCCGCGGACTC-3′ and 5′-AGAAGCGGAACCTCAAATTG-3′, respectively. Pools were then sorted into 96 wells using flow cytometry. Knock-in clones were initially screened by PCR and subsequently confirmed by DNA sequencing.

### Immunoprecipitation (IP)

Cells were lysed by cold lysis buffer. The lysates were centrifuged at 10,000 rpm for 10 min at 4°C. The supernatant was incubated with anti-DYKDDDDK G1 Affinity Resin (GenScript; L00432-10) for 3 h at 4°C. The beads were then washed three times with lysis buffer, followed by the addition of loading buffer containing 10% SDS. The mixture was boiled for 10 min and analyzed by SDS-PAGE.

### Fluorescence recovery after photobleaching (FRAP)

FRAP experiments were performed using a Zeiss Cell Discoverer 7 confocal microscope equipped with a duckweed lens. Cells were imaged for a total of 200 seconds at a maximum rate of 1 frame per second. Photobleaching was carried out with 100% laser power (488 nm wavelength of an argon laser). After 3 pre-bleaching images, the fluorescence signal was bleached from a region of interest (2 μm × 2 μm). The signal from the bleached ROI was then measured for 200 seconds. The photobleached area was imaged every 500 ms. Finally, the fluorescence curves were analyzed by EasyFRAP software (https://easyfrap.vmnet.upatras.gr/).

### Structural prediction

PDB files of SRSF1 (6HPJ) and DDX18 (8FKV) were obtained from the AlphaFold Protein Structure Database (https://alphafold.com/) and then use AlphaFold2 to simulate protein-protein interactions. Simulation results and interaction sites were visualized by PyMOL.

### Protein sequence analysis

Human SRSF1 sequence was downloaded from UniProt (accession No. Q07955). The distribution of charge is defined as the sum of the charges of charged amino acids (R and K, +1; D and E, -1) within an artificial defined window range (window size = 20). The charge plot was generated using an in-house generated Python code.

### EU incorporation assay

U-2 OS cells were incubated with 500 µM 5-ethynyluridine (EU; Beyotime ST2055) for 20 min. Cells were washed with PBS for 5 minutes before being fixed by 4% PFA for 30 min. EU was visualized using Apollo 567/643 (C10371-1/10371-2) according to the manufacturer’s manuals.

### Changing intracellular pH

The buffers used to change the intracellular pH consist of 10 mM PIPES-KOH, 100 mM NaCl, 300 mM sucrose, 1.0 mM EGTA, 1 mM MgCl_2_, 1 mM DTT, and 0.5% Triton X-100. The buffer was freshly prepared on the day of the experiment and stored on ice. Cells were seeded in 3.5 cm plates with glass bottom. Right before the experiments, cells were washed three times with ice-cold PBS, and then were incubated with buffer with the desired pH for 5 min on ice. The cells were then immediately fixed with 4% PFA and processed for standard immunofluorescence.

### Measuring nucleolar pH in cells

To obtain a ratiometric measurement of intracellular pH, we used a fluorescent dye BCECF-AM (Beyotime Institute of Biotechnology, Shanghai, China) as described in previous studies^17,18^. The dye shows pH-sensitive emission when excited at 490 nm, while exhibiting stable emission profile when excited by 440 nm wavelength. Emission was recorded after excitation the dye at wavelengths of 405 and 488 nm, respectively, with a Zeiss LSM980 confocal laser microscope system (Zeiss).

A standard curve for BCECF-AM dye (2 μM) was first generated to analyze intracellular pH. Calibrated PBS with pH values of 6.0, 6.5, 7.0, 7.5 and 8.0 was prepared. 10 μM Nigericin (Shanghai Yuanye Bio-Technology Co. Ltd, Shanghai, China) was added to the calibration buffer 5 min before pH measurement. After calculating the fluorescence ratio of 488/405, we obtained the standard curve of pH. For each experiment to measure nucleolar pH, KI cells with endogenous nucleolar markers tagged with fluorescent proteins were used. The images were first masked using the respective nucleolar markers. The emission ratio at the wavelengths ∼490 nm and ∼440 nm within the nucleolar masks was determined for each individual cell. Finally, we calculated the pH value of the nucleoli of each cell by fitting the ratiometric measurement into the standard curve. The data were pooled for each experimental condition and presented as median ± 95% confidence interval. Each experiment was repeated at least 2 independent times with similar observations.

### Fluorescence in situ hybridization (FISH)

For in situ hybridization, probes were synthesized, labeled with Cy3 (Shenggong, Shanghai) and hybridized as previously described^15^. The RNA probe for 5’-ETS used as previously reported^7^: 5’-CTCTCAGATCGCTAGAGAAGGCTTTTCTCACCGAGGGTGGGTCACACTCC-3’. Cells were washed with PBS and fixed with 4% PFA for 15 min, then permeabilized with 70% ethanol at 4°C for overnight. After washed with the washing buffer (2× SSC, 50% formamide) for 5 min at RT, cells were incubated in the hybridization solution (2× SSC, 200nM vanadyl-ribonucleoside complex, 50% formamide, 0.02% BSA), containing 100 nM RNA FISH probes at 37°C for 4 h. The coverslips were mounted with mounting medium.

### Spectrum analysis

293T cells transfected with pc3.3-Flag-SRSF1 were treated with UV (30 J/m^2^) and recovered for 7 h. Cleared cell lysates were incubated with anti-DYKDDDDK G1 Affinity Resin (GenScript; L00432-10) for 3 h at 4°C. The beads were then eluted with 3x Flag peptide and separated by SDS-PAGE. Trypsin-digested peptides were analyzed using LC-MS/MS (Bruker). Mass spectra were processed using Perseus.

### Statistics and Quantification

The data used in this study were expressed as mean ± SD or mean ± SEM in three replicates. Statistical significance was assessed through Student’s t test or ANOVA, as appropriate. GraphPad Prism 8 software was used for data analysis and p < 0.05 was considered statistically significant. Representative microscopic or western blot images were from at least three independent experiments. The statistical significance and sample size of all the graphs are detailed in the figure legends.

## Acknowledgements

All the imaging data were acquired in the Core Facility of Biomedical Sciences, Xiamen University with the help from Funiu Qin. This work was supported by the National Natural Science Foundation of China (32470726), the Natural Science Foundation of Fujian Province, China (2023J01024, and 2024J010004), the Fundamental Research Funds for the Central Universities.

## Author contributions

B.W. and S.M. designed the experiments. S.S., J.G., X.Z., S.Y., C.H., and B.W. performed experiments and/or analyzed the data. B.W. wrote the manuscript.

## Data availability

Mass spectrometry datasets have been deposited to the ProteomeXchange Consortium with the dataset identifier PXD062285. Source data are provided with this paper. All other data supporting this study are available from the corresponding author upon reasonable request.

## Figure legends

**Figure S1 (related to Figure 1).**
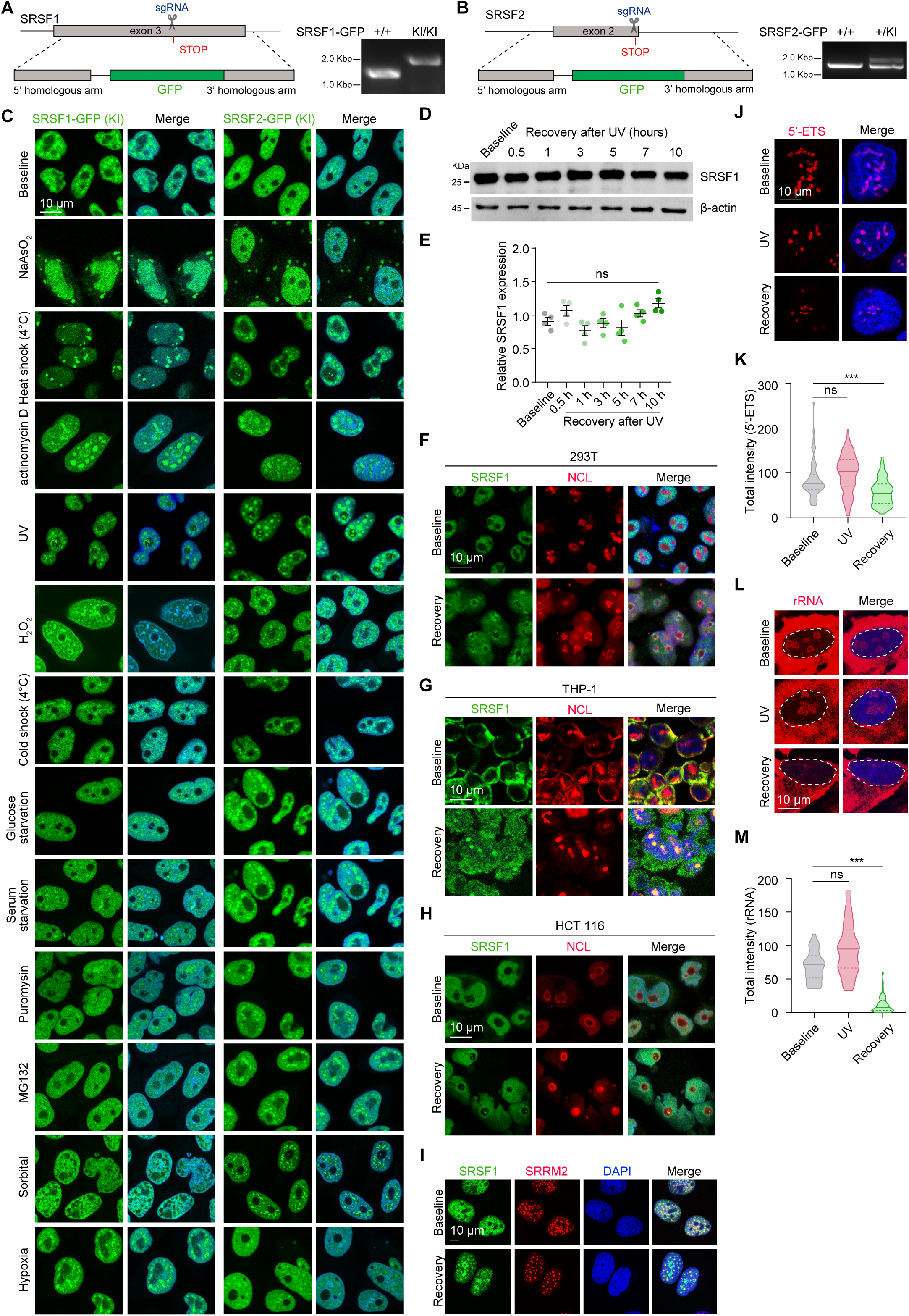
SRSF1 is recruited into nucleolus under specific stressed conditions. (**A**, **B**) Schematic illustration of SRSF1-GFP KI (A) and SRSF2-GFP KI (right), and screening by PCR. (**C**) Representative confocal images of SRSF1-GFP and SRSF2-GFP KI HeLa cells treated with the indicated stressors. (**D**, **E**) Lysates prepared from WT U-2 OS cells subjected to the indicated treatment were analyzed by immunoblotting. The relative SRSF1 expression levels were measured (E). (**F**-**H**) WT 293T (F), THP-1 (G), or HCT 116 (H) cells were treated with UV and recovered for 7 h in normal growth condition. The cells were then immunostained with antibodies against SRSF1 and NCL. (**I**) SRSF1-GFP KI U-2 OS cells were treated with UV and recovered in normal growth condition for 7 h. The cells were immunostained with SRRM2 antibody. (**J**, **K**) U-2 OS cells treated with UV and recovered in normal growth condition for 7 h were subjected to FISH using a probe against 5’-ETS. The total intensity of 5’-ETS in each cell was determined. (**L**, **M**) U-2 OS cells treated with UV and recovered in normal growth condition for 7 h were subjected to immunostaining using an rRNA antibody. The total intensity of rRNA in the nucleolus was determined. For (**E**), data are shown as mean ± SEM. n = 3 independent experiments. ns, not significant by one-way ANOVA. For (**K**), and (**M**), data are shown as mean ± SD. ns, not significant; **p < 0.01 by one-way ANOVA.

**Figure S2 (related to Figure 2).**
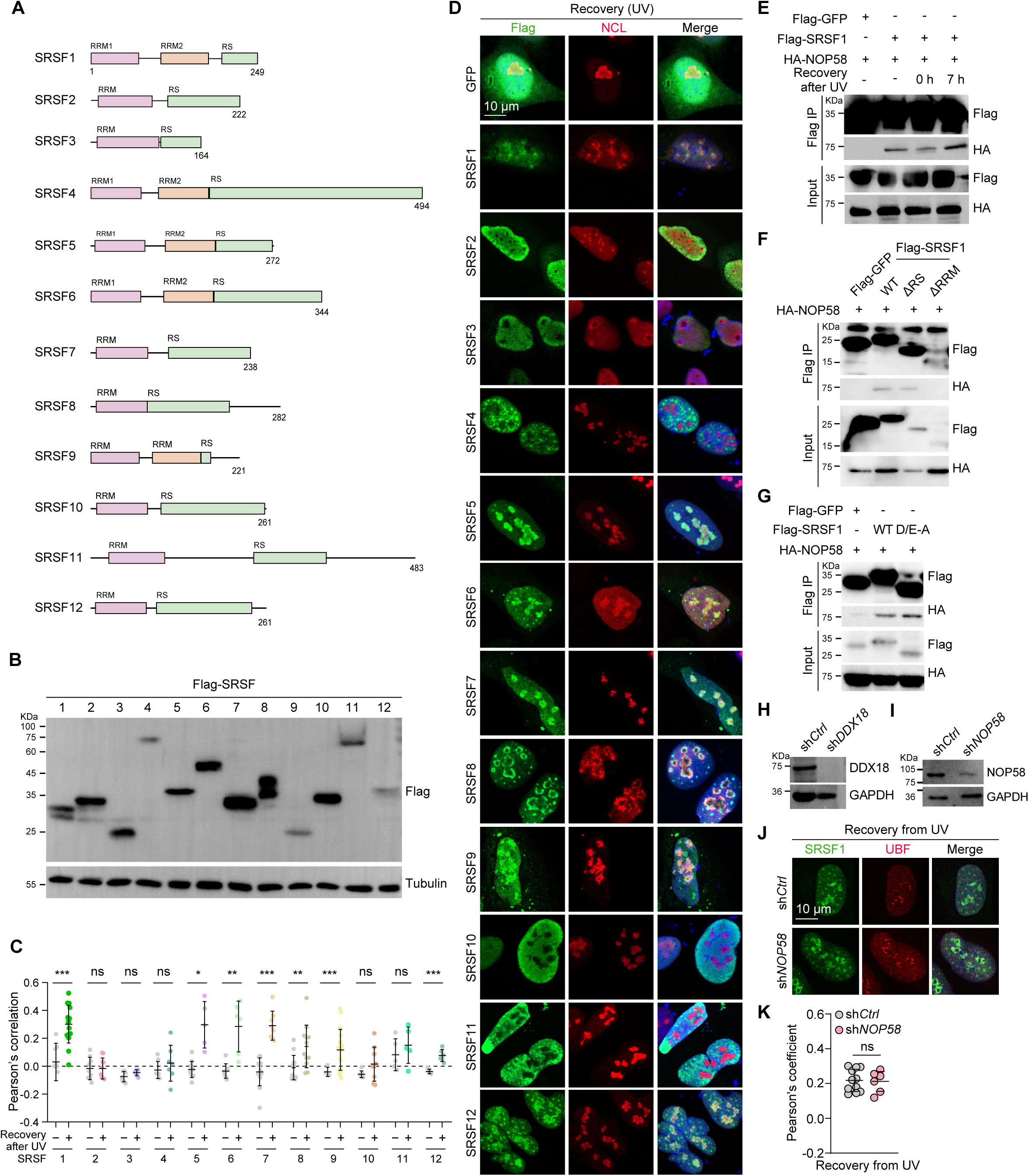
SRSF1 localization to the nucleolus is dependent on the molecular interaction with DDX18 through an acidic charge block. (**A**) Schematic illustration of the domains of the SRSF family members. (**B**) 293T cells were transfected with the indicated constructs and analyzed by immunoblotting. (**C**, **D**) WT U-2 OS cells transfected with the indicated Flag-tagged SRSF proteins were stressed by UV and allowed to recover for 7 h. The cells were immunostained with the indicated antibodies (C). Pearson’s correlation of SRSF proteins and NCL were quantified (D). (**E**) 293T cells were transfected with the indicated constructs and subjected to Flag immunoprecipitation. (**F**) 293T cells transfected with the indicated constructs were stressed with UV and recovered for 7 h before being subjected to Flag immunoprecipitation. (**G**) 293T cells were transfected with the indicated constructs and subjected to Flag immunoprecipitation. (**H**, **I**) Lysates prepared from WT U-2 OS cells infected with shRNA against *Ctrl*, *DDX18* (H), or *NOP58* (I) were analyzed by immunoblotting using the indicated antibodies. (**J**, **K**) WT U-2 OS cells infected with shRNA against *Ctrl* or *NOP58* were stressed with UV followed by recovery in normal growth condition for 7 h. The cells were then processed for immunostaining with the indicated antibodies (J). Pearson’s correlation of SRSF1 and UBF were quantified (K). Data are shown as mean ± SD. ns, not significant; *p < 0.05, **p < 0.01; ***p < 0.001 by unpaired Student’s t test.

**Figure S3 (related to Figure 3).**
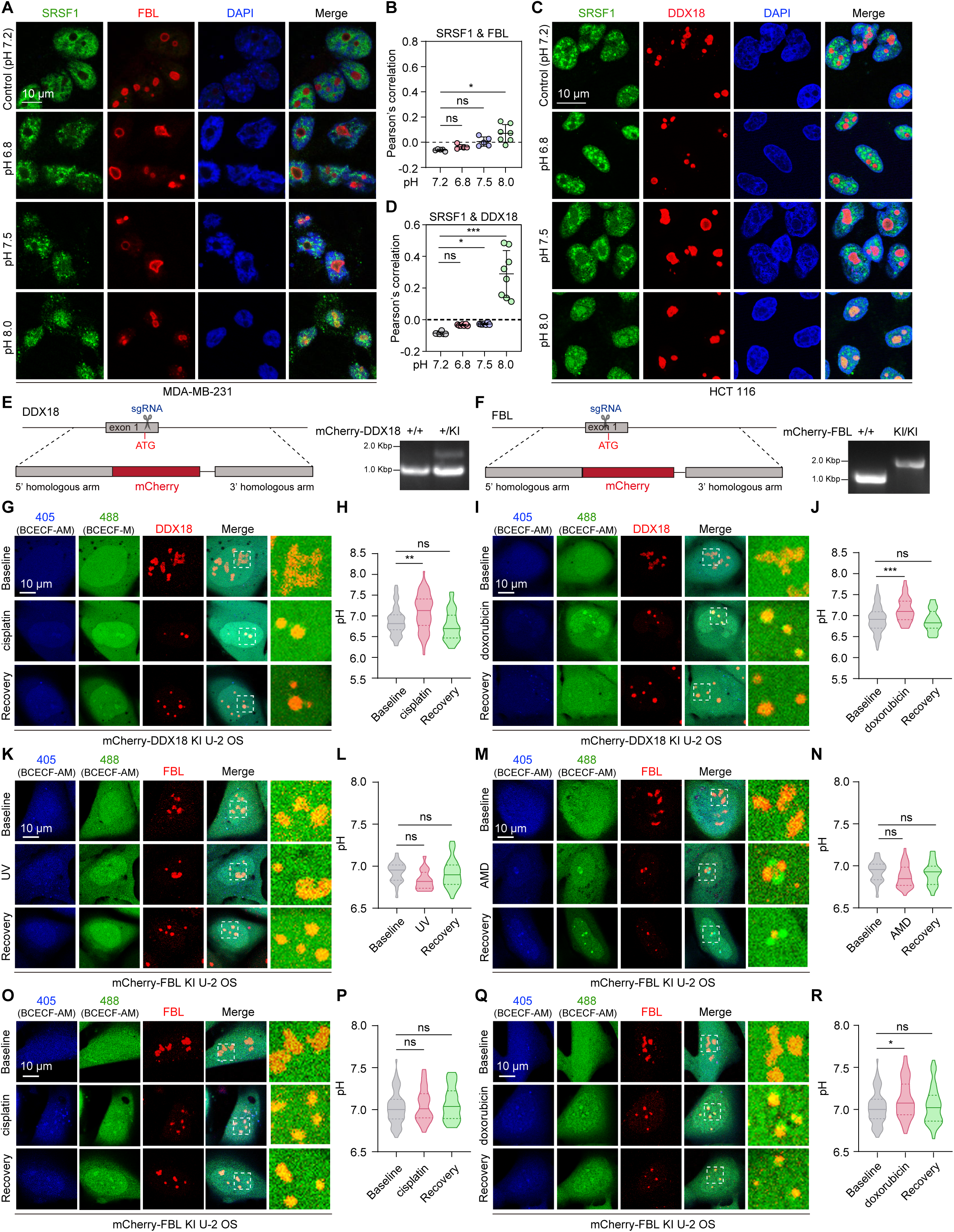
SRSF1 senses and maintains nucleolar pH. (**A**-**D**) WT MDA-MB-231 (A, B) or HCT 116 (C, D) cells were cultured in medium at varying pH. The cells were immunostained with the indicated antibodies. Colocalization of SRSF1 with the nucleolus was quantitatively assessed by Pearson’s correlation. Data are shown as mean ± SD. ns, not significant; *p < 0.05; ***p < 0.001 by one-way ANOVA. (**E**, **F**) Schematic illustration of mCherry-DDX18 KI (E) and mCherry-DDX18 KI (F), and screening by PCR. (**G**-**N**) Live cell imaging of mCherry-FBL KI U-2 OS cells being treated with either UV (G, H), AMD (I, J), cisplatin (K, L), or doxorubicin (M, N) and recovering in normal growth condition with a pH indicator BCECF-AM. pH was quantified by calculating the ratio of the fluorescence intensity of 488/405 nm in the nucleoli. ns, not significant; *p < 0.05 by one-way ANOVA. (**O**-**R**) Live cell imaging of mCherry-DDX18 KI U-2 OS being treated cisplatin (O, P), or doxorubicin (Q, R) and recovered in normal growth condition with BCECF-AM. ns, not significant; **p < 0.01; ***p < 0.001 by one-way ANOVA.

**Figure S4 (related to Figure 4).**
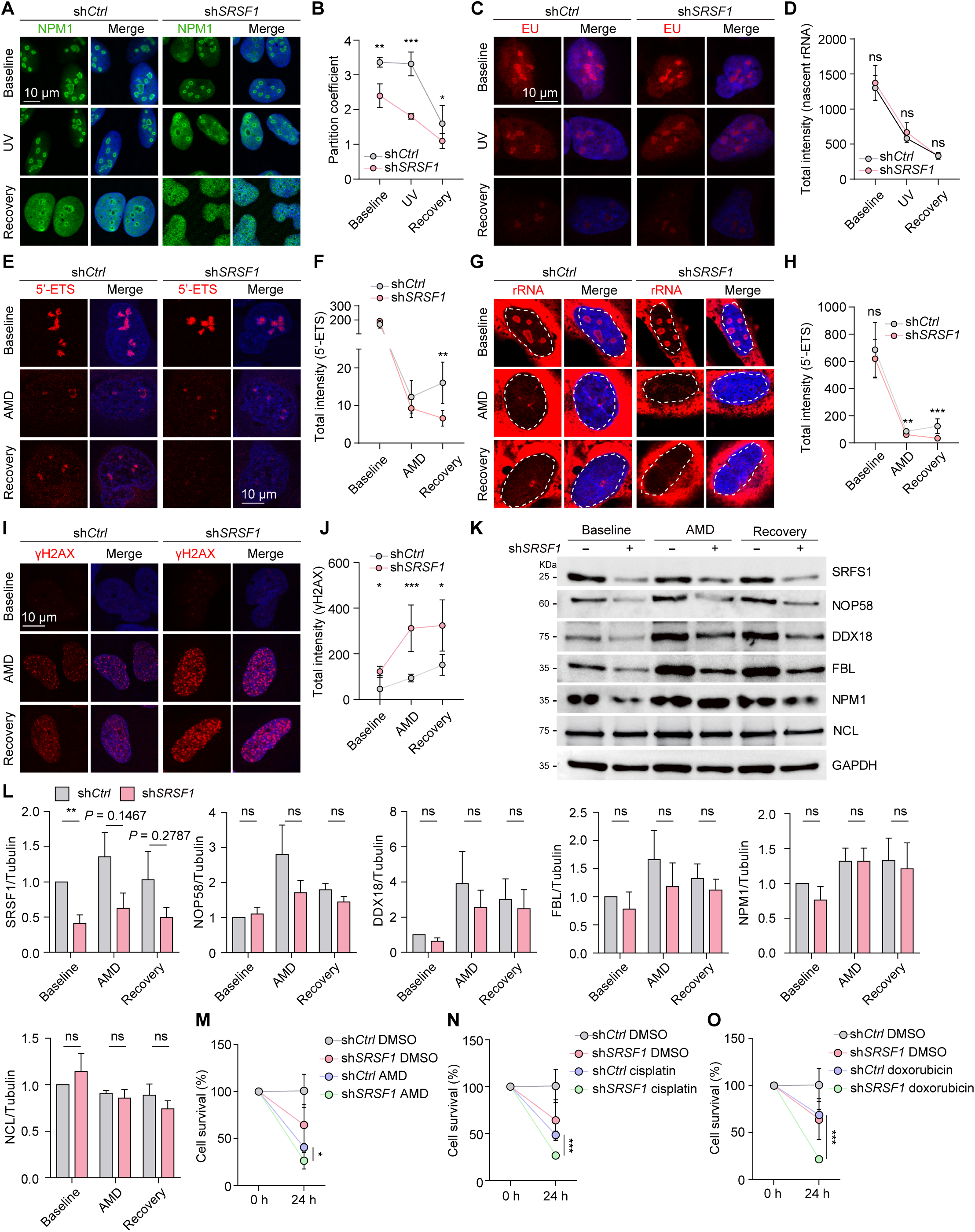
SRSF1 maintains nucleolar integrity and function in times of stress. (**A-D**) WT U-2 OS cells transduced with shRNA against *Ctrl* or *SRSF1* were treated with UV, and allowed to recover in normal growth condition for 7 h. The cells were immunostained with a NPM1 antibody. The partition of NPM1 into the nucleolus was quantified (B). Nascent RNA was labeled by EU incorporation (C, D). (**E-I**) WT U-2 OS cells transduced with shRNA against *Ctrl* or *SRSF1* were treated with AMD and recovered for 4 h. The cells were subjected to either FISH using a probe for 5’-ETS (E, F), immunostaining using an antibody against rRNA (G, H), or γH2AX (I, J). The total intensity of 5’-ETS (F), rRNA (H), and γH2AX (J) were quantified, respectively. (**K**, **L**) WT U-2 OS cells transduced with shRNA against *Ctrl* or *SRSF1* were treated with UV, and allowed to recover in normal growth condition for 7 h. The cells were lysed and analyzed by immunoblotting using the indicated antibodies. The relative band intensities were quantified by densitometry (L). (**M**-**O**) WT U-2 OS cells transduced with shRNA against *Ctrl* or *SRSF1* were treated with AMD (M), cisplatin (N) or doxorubicin (O). 24 h after the treatment, cell survival was determined by CCK8 assay. For (**B**), (**D**), (**F**), (**H**), and (**J**), data are shown as median ± 95% confidence interval. ns, not significant; *p < 0.05, **p < 0.01, ***p < 0.001 by unpaired Student’s t test. For (**L**), data are shown as mean ± SEM. n = 3 independent experiments. ns, not significant; **p < 0.01 by unpaired Student’s t test. For (**M**), (**N**), and (**O**), data are shown as mean ± SD. ns, not significant; *p < 0.05; ***p < 0.001 by unpaired Student’s t test.

**Figure S5 (related to Figure 5).**
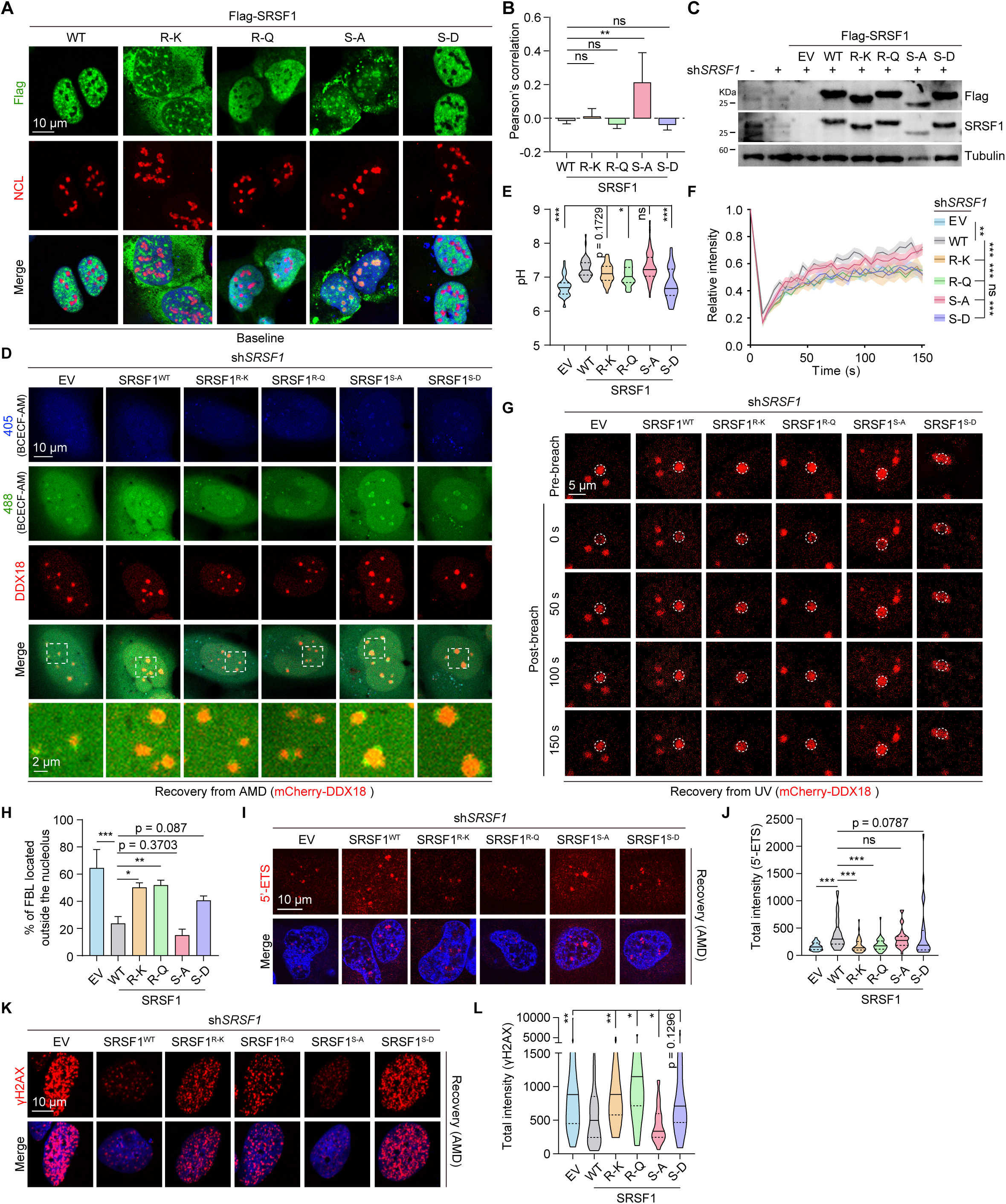
SRSF1 modulates nucleolar pH and function via its alkaline RS domain. (**A**, **B**) U-2 OS cells expressing the indicated SRSF1 variants were immunostained with the indicated antibodies. Pearson’s correlation of SRSF1 and NCL were quantified (B). Data are shown as mean ± SD. (**C**) Lysates from the indicated cell lines were analyzed by immunoblotting. (**D**, **E**) The indicated cell lines were treated with AMD. Nucleolar pH was measured by live cell imaging. (**F**, **G**) The indicated cells were treated with UV and recovered for 7 h. The cells were subjected to FRAP. The fluorescence intensity of mCherry-DDX18 was monitored over time and quantified (F). Data are shown as mean ± SD. n = 15. *p < 0.05 by two-way ANOVA. (**H**) The indicated cell lines were treated with AMD and recovered for 4 h. The cells were immunostained with antibodies against DDX18 and FBL. Percentage of nucleoli with FBL dislocated to the exterior were quantified. Data are shown as mean ± SEM. n = 3 independent experiments. (**I**-**L**) The indicated cells were treated with AMD and recovered for 4 h. The cells were processed for either FISH using a probe for 5’-ETS (I, J), or immunostaining using an antibody against γH2AX (K, L). The total intensity of 5’-ETS (J) and γH2AX (L) were quantified, respectively. For (**B**), (**E**), (**H**), (**J**), and (**L**), ns, not significant; *p < 0.05; **p < 0.01; ***p < 0.001 by one-way ANOVA. For (**F**), ns, not significant; **p < 0.01; ***p < 0.001 by two-way ANOVA.

**Figure S6 (related to Figure 6).**
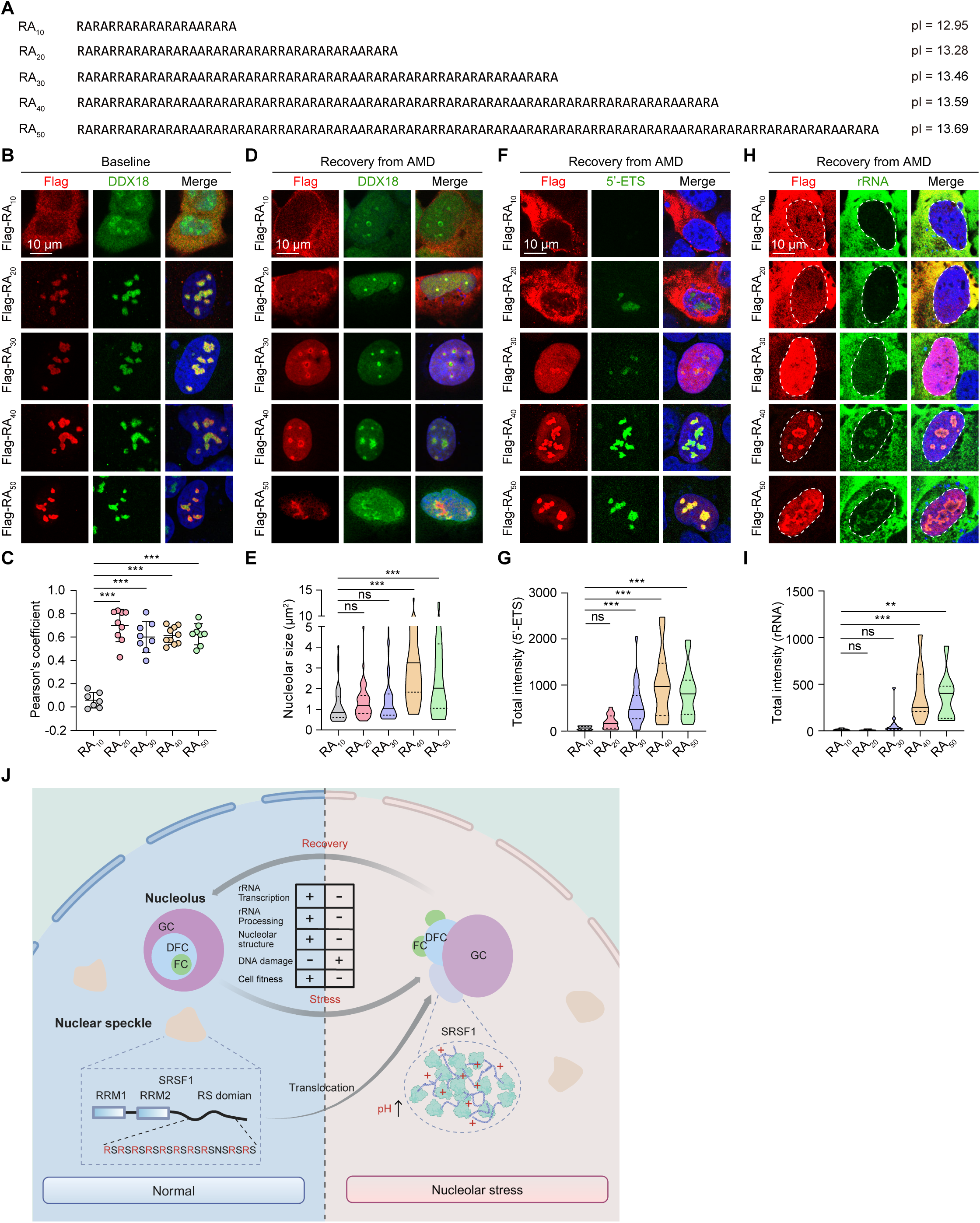
Synthetic arginine-rich alkaline dipeptide protects nucleolus from stress that disrupt nucleolar pH and function. (**A**) The linear sequences of synthetic dipeptides. The corresponding pI was shown on the right. (**B**, **C**) U-2 OS cells expressing RA dipeptides of different lengths were immunostained with the indicated antibodies. Pearson’s correlation between DDX18 and different dipeptides was quantified (C). Data are shown as mean ± SD. ***p < 0.001 by one-way ANOVA. (**D**-**I**) U-2 OS cells expressing RA dipeptide were first treated with AMD, followed by recovery in fresh medium for 4 h. The cells were further processed for FISH using a probe against 5’-ETS or immunostaining using the indicated antibodies. The nucleolar size (E), total intensity of 5’-ETS (G), and rRNA (I) were quantified, respectively. ns, not significant; **p < 0.01; ***p < 0.001 by one-way ANOVA. (**J**) Proposed model of this study.

